# Microglial engulfing of glutamatergic inputs and diminished excitability of D1 medium spiny neurons of the nucleus accumbens by peripheral inflammatory insult

**DOI:** 10.1101/2024.12.02.625901

**Authors:** Shingo Nakajima, Anita Kabahizi, Solal Aubailly, Samar Naili-Douaouda, Akari Tomita, Anthony Bosson, Ciaran Murphy-Royal, Stephanie Fulton

## Abstract

Neuroimmune responses to systemic inflammation can generate anxiodepressive behaviors and psychomotor slowing. The nucleus accumbens (NAc) is a neural hub encoding mood and motivation that contributes to the locomotor and mood dampening effects of neuroinflammation. Dopamine receptor 1 mediums spiny neurons (D1-MSNs) of the NAc regulate locomotor activity, motivated behavior and emotional states and are modulated by microglial reactivity. Here, we evaluated sickness- and anxiety-like behaviors along with D1-MSN activity and microglial responses in the NAc to systemic lipopolysaccharide (LPS) administration. LPS stimulated anxiety-like behavior, blunted locomotion and reduced cFos expression in D1-MSNs of the NAc core and shell of male mice. These effects associated with reduced excitatory inputs (EPSCs) onto D1-MSNs as measured by whole cell patch-clamp. To determine if microglia contribute to changes in MSN activity, Ca^2+^ imaging of primary cultures containing NAc neurons with or without primary NAc microglia was performed. The presence of microglia decreased the activity of LPS-treated MSNs in response to dopamine and glutamate application and LPS stimulated microglial phagocytosis of MSN processes in co-cultures. Immunohistochemical analyses revealed that *in vivo* LPS treatment enhanced morphological indices of microglia reactivity and engulfment of vesicular glutamate transporter 1 (VGLUT1) inputs in the NAc. Our results suggest that LPS *reduces locomotion* and stimulates anxiety via increasing microglia engulfment of excitatory inputs onto D1-MSNs, and highlight changes in NAc microglia phagocytic activity in the behavioral consequences of neuroinflammation.

## INTRODUCTION

Neuroinflammation is increasingly recognized as a source of mood and motivational disturbances in conditions of obesity and associated metabolic endotoxemia (1, 2, 3). Gut-derived endotoxins like lipopolysaccharide (LPS), which are upregulated in obesity, stimulate peripheral immune signals that are relayed to the brain via autonomic afferents and circulating cytokines and chemokines (4, 5, 6). Ensuing neuroimmune mechanisms are well-implicated in sickness behaviors that promote adaptive recovery in rodents (7, 8, 9, 10) and humans (11). As resident immune cells of the brain, microglia are central to neuroinflammatory responses and essential for regulating neuron connectivity and health. Microglia engulf and remove synaptic elements, a process known as phagocytosis, and thereby refine neural circuits and modulate behavior (12, 13). The nucleus accumbens (NAc), with its anatomically distinguished core and shell sub-regions, is a key brain region encoding motivation and mood (14). The NAc contains dopaminoceptive GABAergic medium spiny neurons (MSNs) expressing D1 (D1-MSNs) or D2 (D2-MSNs) receptors that contribute to effortful behavior, reinforcement and emotional states, (15, 16, 17). Consistent with the NAc as a neural hub for behavioral regulation (18, 19), D1- and D2-MSNs integrate excitatory inputs from the basolateral amygdala (BLA) and prefrontal cortex (PFC) in the NAc core (20) and from the PFC, ventral hippocampus, and medio-dorsal thalamus in the NAc shell (21, 22). The traditional view of D1-MSNs and D2-MSNs in the striatum encode opposing valences, with D1-MSN activation producing positive reinforcement and D2-MSN activation leading to aversion and negative reinforcement. Generally, D1 neuron activation promotes stress resilience and antidepressant-like effects in chronic social defeat stress models (23). In contrast, some studies suggest that D2 neuron activation may promote depressive-like behaviors (23). However, recent research has revealed that behavioral outcomes are more nuanced and depend on specific projection targets. There is evidence linking both D1- and D2-MSN to anxiety-like behavior (15, 24). These findings highlight the complexity of striatal outputs and their diverse effects on behavior and emotion regulation.

Systemic inflammation caused by endotoxemia and chronic obesogenic diets induces anxiodepressive behaviors along with increased gliosis and inflammatory markers in the NAc (25, 26). Microglia play a significant role in regulating synaptic plasticity through the purinergic signaling and phagocytosis (27, 28, 29). Several lines of evidence indicate that peripheral inflammatory responses activate microglial phagocytosis to engulf neuronal fibers (28, 30, 31); however, it is unclear how NAc microglia interact with MSNs during neuroinflammation. To address this question, we assessed anxiodepressive behavior, MSN activation and MSN-microglia interactions in response to LPS using immunohistochemistry, whole-cell recordings and live cell imaging.

## MATERIALS AND METHODS

### Animals

All procedures involving the use of animals were approved by the CRCHUM Animal Care Committee in accordance with Canadian Council on Animal Care guidelines. Male C57Bl/6 mice (Jackson Laboratory, Bar Harbor, ME, USA) and D1R^Cre^ mice (C57Bl/6 background; in-house colony) were housed in a reverse 12h light-dark cycle (lights off at 10h) with ad libitum access to water and food. Colony-derived heterozygous D1R^Cre^ mice and wildtype littermates were weaned at postnatal day 21-23 and genotyping of transgenic mice was performed by PCR analysis from ear biopsies.

### Procedures for behavioral assessment

To assess the behavioral effects of acute inflammation, mice were intraperitoneally injected with lipopolysaccharide (LPS; 0.83 mg/kg, serotype 0127:B8, Sigma-Aldrich, St. Louis, MO, USA) or vehicle (saline). 12 hours post-injection, mice underwent behavioral testing in the following sequence: elevated plus maze (EPM), open field test (OFT), light/dark box (LDB), and three chamber social interaction test (3CT) for the assessment of anxiety-like and social behaviors. The LPS dose was chosen based on a report showing it is the minimal effective dose to induce anxiety- and depressive-like behavior. Control mice received an IP injection of vehicle (endotoxin-free saline solution). All behavior tests were captured by an overhead video camera connected with Ethovision XT software (Noldus, Leesburg, VA, USA) for a period of five minutes.

### Elevated-plus maze

The EPM was performed as previously reported (9). Briefly, each mouse was placed in the center of the maze facing open arm. Total distance traveled and time spent in each arm were analyzed. The percentage of time spent in open arm was calculated as the ratio of open arm duration and total spent time in each arm.

### Open field

Mice were placed into a corner of an open field apparatus (40 cm X 40 cm X 27 cm). The center zone was set 13.3 cm x 13.3 cm in the arena. Total traveled distance and time spent in center were analyzed. The percentage of time spent in center was calculated as the ratio of center zone duration and total test time.

### Light/dark box

The apparatus consisted of an illuminated compartment made of transparent plastic and a dark compartment made of black plastic, covered by a lid (both 13.7 cm X 13.7 cm X 20.3 cm). The two boxes were separated by a partition wall, with an opening at the bottom to allow the animal to pass freely between compartments. Total traveled distance in light zone and time spent in in each zone were analyzed. The percentage of time spent in light zone was calculated as the ratio of light zone duration and total test time.

### Three-chamber social interaction

The three-chamber social interaction test (3CT) is used to assess general sociability. The rectangular apparatus (40 cm X 60 cm X 23 cm) contains three connected compartments divided by opaque Plexiglas walls. An unfamiliar male mouse was put in a small wire cage on one side of the chamber as the stimulus mouse, whereas an empty wire cage put on another side. Total distance and contact time with both the stimulus mouse and empty cage were analyzed. The contact time was manually scored to analyze the duration in each contract.

### Immunohistochemistry

Mice were trans-cardially perfused with phosphate buffered saline (PBS) followed by 10% formalin 30-45 minutes after 3CT behavior testing. Dissected brains were post-fixed in 10% formalin overnight followed by 20% sucrose until brains sunk. Coronal slices were obtainedat 30 µm intervals with microtome (Leica). After washing with PBS and blocking (2 hours in 3% NGS, 0,1% Triton, in PBS), slices were incubated with primary antibodies for anti-Cre (1:500, Guinea pig, Synaptic systems), anti-cFos (1:500, Rabbit, Cell Signaling), anti-Iba1 (1:1000, Rabbit, Fujifilm-WAKO), anti-CD68 (1:1000, rat, Thermo fisher), anti-VGLUT1, 1:1000, Guinea pig, Sigma Aldrich) in the blocking solution overnight at 4°C. The brain slices were washed several times with PBS followed by secondary antibody incubation (Anti-guinea pig IgG Alexa 488, Anti-rabbit IgG Alexa 488, Anti-rabbit IgG Alexa 568, and/or Anti-rat IgG Alexa 647, Invitrogen) for 2h at room temperature. Brain sections were mounted with vectashield contains DAPI (Vector Labs) and imaged with Axio Imager A2 Zeiss fluorescent microscope (Carl Zeiss AG). 15-20 Z-stack images were captured with a step size of 1 µm and stacked in ImageJ2/FIJI (32). Cre and c-Fos colocalization was analyzed with the “colocalization” function. Microglia soma size was measured after converting to binary image with equivalent threshold and then region of interest (ROI) was created from each microglia. All vesicle of CD68 and VGLUT1 in the ROI were measure with analyze particle function.

### Stereotaxic Surgery

D1R^Cre^ mice were anesthetized with isoflurane and placed into a mouse ultraprecise stereotaxic instrument (Kopf, Inc.) with Bregma and Lambda in the same horizontal plane. To visualize cell bodies for patching D1-MSNs, AAV5-EF1a-DIO-eYFP (Addgene plasmid # 27056; http://n2t.net/addgene:27056; RRID: Addgene_27056) was bilaterally (500 nl) injected into the NAc of D1R^Cre^ mice (AP:+1.5 mm. ML:±1.3 mm, DV: −4.3 mm, relative to bregma and skull surface) with using a 2 μL NeuroSyringe at (Hamilton, Reno, NV, USA). AAV5 titers were 3.7– 6 × 10^12^ viral molecules/ml. For whole-cell patch-clamp experiments, D1R-cre reporter mice underwent viral stereotaxic surgery at 8 weeks of age and thereafter were singly housed. Mice were used for experimentation at 2-3 weeks following surgery.

### Brain slice preparation for electrophysiology

Animals were injected saline or LPS (0.83 mg/kg body weight) 12 h before isoflurane anesthesia and trans-cardially perfused with ice-cold, oxygenated (95% O_2_/5% CO_2_) N-methyl-d-glutamine (NMDG) containing recovery solution (in mM: 119.9 NMDG, 2.5 KCl, 25 NaHCO_3_, 1 CaCl_2_-2H_2_O, 2 MgCl_2_-6H_2_O, 1.4 NaH_2_PO_4_-H_2_O, 20 Glucose). Brains were rapidly dissected and washed in ice cold, oxygenated NMDG. 300 µm coronal slices were collected in ice cold, oxygenated NMDG solution with Leica VT 1200S vibratome. Slices were transferred to 32° oxygenated NMDG solution in a recovery chamber for 12 min before being stored at room temperature oxygenated artificial cerebrospinal fluid (ACSF) (in mM: 130 NaCl, 2.5 KCl, 1.25 NaH_2_PO_4_-H_2_O, 1.3 MgSO_4_, 2 CaCl_2_, 10 Glucose, 26 NaHCO_3_, 287-295 mOsm). Slices rested for at least 1 h prior to recording. For electrophysiological recording, slices were moved to a recording stage perfused with oxygenated ACSF at a rate of 200 ml/hr at 30°C using an inline heater.

### Whole-cell patch-clamp recordings

For excitatory current whole-cell recordings, the pipette solution included: in mM, 130 K-Gluconate, 10 Hepes, 5 KCl, 5 NaCl, 4 ATP-Mg, 0.3 GTP-Na, 10 Na_2_-Creatine-PO_4_. For inhibitory current whole-cell recordings, the pipette solution included: in mM, 130 CsCl, 10 NaCl, 10 Hepes, 1 EGTA, 0.1 CaCl_2_, 10 PCr, 4 ATP-Mg, 0.4 GTP-Na, 5 Qx-314 (Lidocaine). Electrophysiological signals were recorded using an Axopatch 700B amplifier (Molecular Devices), low-pass filtered at 2–5 kHz, and analyzed offline on a PC with pCLAMP programs (Molecular Devices). Excitatory (EPSC) and inhibitory (IPSC) postsynaptic currents were measured by whole-cell voltage clamp recordings, and membrane potential rates were measured by whole-cell current clamp recordings from D1R-expressing neurons in brain slices. Recording electrodes had resistances of 2.5–5 MΩ when filled with the K-gluconate or CsCl internal solution. Only cells with a stable holding current less than +/-100 pA and an access resistance <20 MΩ were included. Furthermore, cells were rejected if the holding current and access resistance changed more than 20% during recording. Solutions containing drugs were perfused for at least 5 minutes. EPSCs were recorded with GABA_A_ receptor blocker picrotoxin (60 µM working solution in <1 % DMSO) and IPSCs were recorded with glutamate receptor blocker DNQX (10 µM working solution in H2O).

### Primary cultured neurons and microglia from ventral striatum

C57BL/6 pups at postnatal day 1-3 were used for primary cultures as previously reported (9, 33). Briefly, the NAc was dissected and minced with a surgical knife. Tissue was treated with an enzymatic solution containing papain (9 U/ml), DNase (200 U/ml), glucose (5 mg/ml), cysteine (0.2 mg/ml), and bovine serum albumin (0.2 mg/ml) for 15 min at 37°C in 5% CO_2_. Following the removal of cell debris by the passing with 70 µm cell strainer and centrifuge, cells were re-suspended with Dulbecco’s modified Eagle’s medium (DMEM) supplemented with 10% heat-inactivated fetal bovine serum (FBS) and 1% of antibiotics (Penicillin G (10,000 U/ml)-Streptomycin Sulfate (10,000 μg/ml) or Neurobasal-A medium containing, 2% B-27 supplement, 1% Glutamax, and 1% antibiotics solution for mixed glial cells or neurons, respectively. To inhibit astrocyte growth, AraC (2 µM) was added at 3 days in vitro (3 DIV). After reaching astrocyte confluence (10-14 days) in mixed glial culture, microglia were isolated from astrocyte layer by the shaking of cultured flask. The number of microglia was counted with hemocytometer followed by seeding of microglia to transwell (5 x 10^4^ cells) or neuronal culture (3 x 10^4^ cells) at 7-8 DIV.

### Live cell calcium imaging in primary cultured neurons

Primary cultured neurons were grown on poly-L-lysine coated 12 mm coverslips on 12-well plates with or without microglia until 11-12 DIV. After LPS treatment (100 ng/ml) for 24 hours, cells were exposed to 3 µM calcium indicator dye Biotracker 609 Red Ca^2+^-AM (Millipore) in Krebs-Hepes Buffer containing (in mM); 135 NaCl_2_, 6 KCl, 2 CaCl_2_, 1.2 MgCl_2_, 10 Glucose, 10 HEPES, at pH 7.3-7.4 for 40-50 min. After flushing dye with fresh Krebs-Hepes buffer, the coverslip was set in the closed perfusion chamber (ALA Scientific instrument inc.). Images were captured with Ex/Em 589 nm/608 nm at 2 hertz with Axio Imager 2 (Zeiss). Data was adjusted according a fluorescence decay curve. The data shows relative change to base line (ΔF/F0). Peak detection was analyzed with ImageJ2/Fiji (32), and the peak as identified by higher intensity above 2SD of baseline.

### Immunocytochemistry

Primary cultured cells on coverslips were fixed with 4% paraformaldehyde followed by several PBS washes. Primary antibodies for MAP2 (Sigma-Aldrich) and IBA-1 (Fujifilm Wako) were applied at 1:1000 dilution overnight. Coverslips were treated with secondary antibodies (Anti-rabbiit IgG Alexa 488 and Anti-mouse IgG Alexa 568, Invitrogen) for 1h at room temperature. Coverslips were mounted with DAPI Vectashield (Vector Labs) and imaged with Axio Imager A2 Zeiss fluorescent microscope (Carl Zeiss AG). MAP2 fragment number and area in microglia were counted with Fiji.

### Statistical analyses

All statistical analyses were conducted using Prism 9 (GraphPad Software, Boston, MA, USA). The data are represented as mean ± SEM. Two-way repeated measures mixed analysis of variance (ANOVA) was done followed by multiple comparisons with Bonferroni’s test for post-hoc analyses on neuron-microglia co-culture experiments. All other data was analyzed using unpaired two-tailed t-test. Differences were considered significant at p<0.05. Asterisks indicate *p<0.05, **p<0.01, ***p<0.001, and ****p<0.0001.

## RESULTS

### Systemic inflammation decreases neuronal activity in NAc D1R neurons

Mice received a systemic LPS injection prior to behavior tests that were followed by cardiac perfusions for immunohistochemical analyses (Figure 1A). LPS reduced anxiety-like behavior in the EPM (Figure 1B) and OFT (Figure 1D) with a tendency to decrease time spent in the light zone of the LDB (Figure 1F). LPS decreased distance travelled in the light zone of LDB (Figure 1G) and in the 3CT (Figure 1H) and produced a trend for decreased distance traveled in the EPM and OFT.

**Figure 1.**
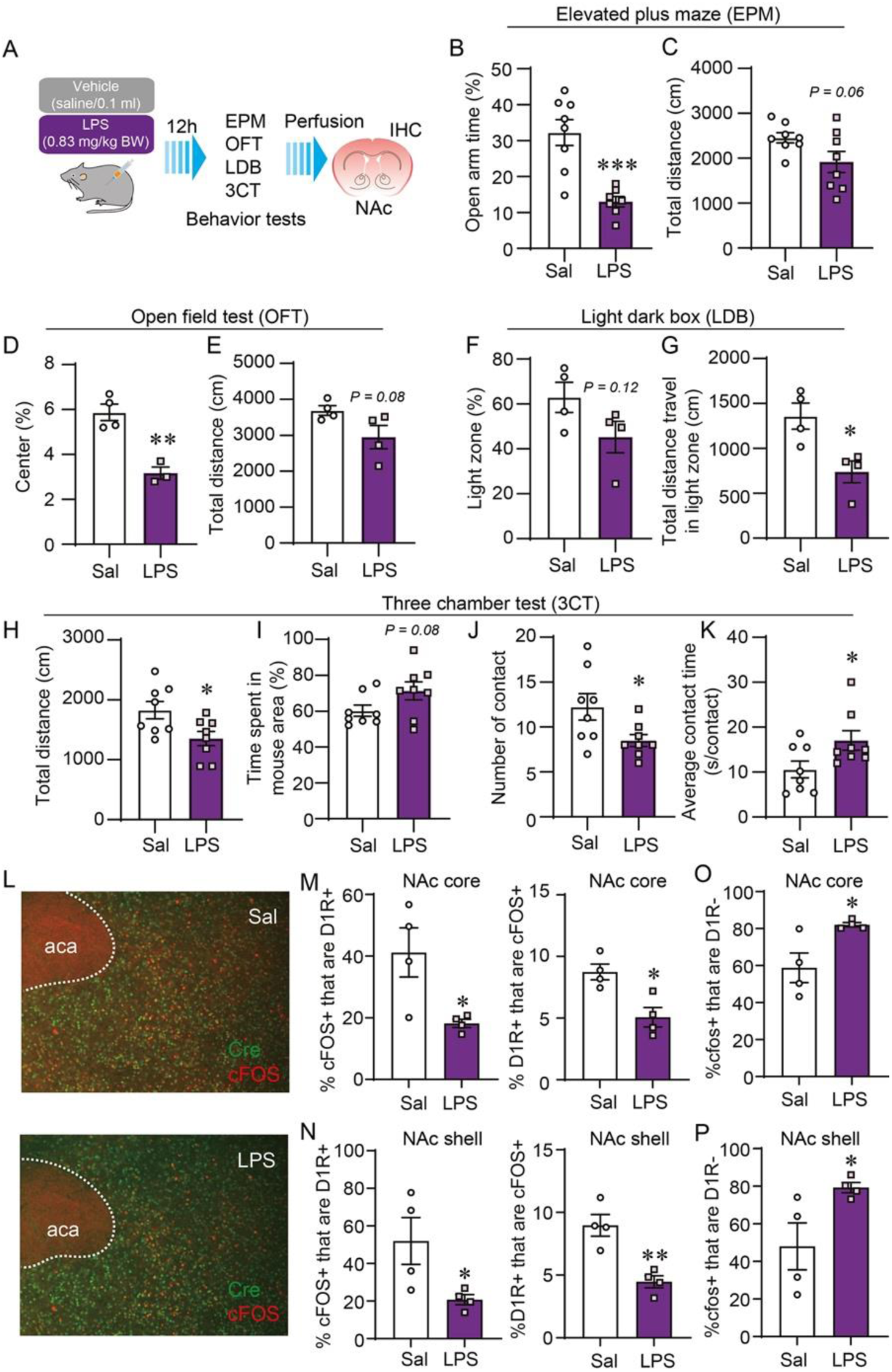
Reduced cFOS-positive D1-MSNs in the NAc core and shell in mice exhibiting anxiety-like behavior caused by LPS injection. (A) The schematic illustration for the test during systemic inflammation (n=4-8 for each treatment). (B) Open arm % and (C) total distance travel in EPM. (D) Center % and (E) total distance travel in OFT. (F) Light zone time % and (G) Total distance travel in light zone in LDB. (H) Total distance travel, (I) time spent in mouse area %, (J) number of contacts, and (K) average contact time in 3CT. (L) Representative images of cFos (red), cre (green) staining in the NAc. (M, N) Plots show % cFOS+ that are D1R positive, % D1R+ that are cfos positive and (O, P) % cfos+ that are D1R negative. All data are expressed as mean ± SEM and analyzed using unpaired *t*-test or non-parametric Mann Whitney test; * p<0.05, **p<0.01, and ***p<0.001.

We next sought to evaluated c-Fos levels in the NAc core and shell to determine changes in neuronal activity. In both subcompartments, the number of c-Fos+ D1-MSNs (Cre+) was decreased by LPS (Figure 1L-N). The number of c-Fos positive cells in Cre-cells (presumably mostly D2-MSN) was increased in the NAc core and medial shell of LPS-treated mice (Figure 1O-P).

### LPS reduces excitatory input onto D1-MSNs

D1R^cre^ mice received bilateral injections of a Cre-dependent adenovirus (Figure 2A) to visualize D1-MSNs for the whole-cell patch-clamp recording (Figure 2B). EPSC frequency, but not amplitude and resting membrane potential (RMP), was reduced in NAc core and medial shell in LPS-treated D1R-cre mice (Figure 2C-J). IPSC frequency was not affected by LPS treatment in (Figure 2K,M). IPSC amplitude was reduced in both the NAc core and shell by LPS (Figure 2K-N).

**Figure 2.**
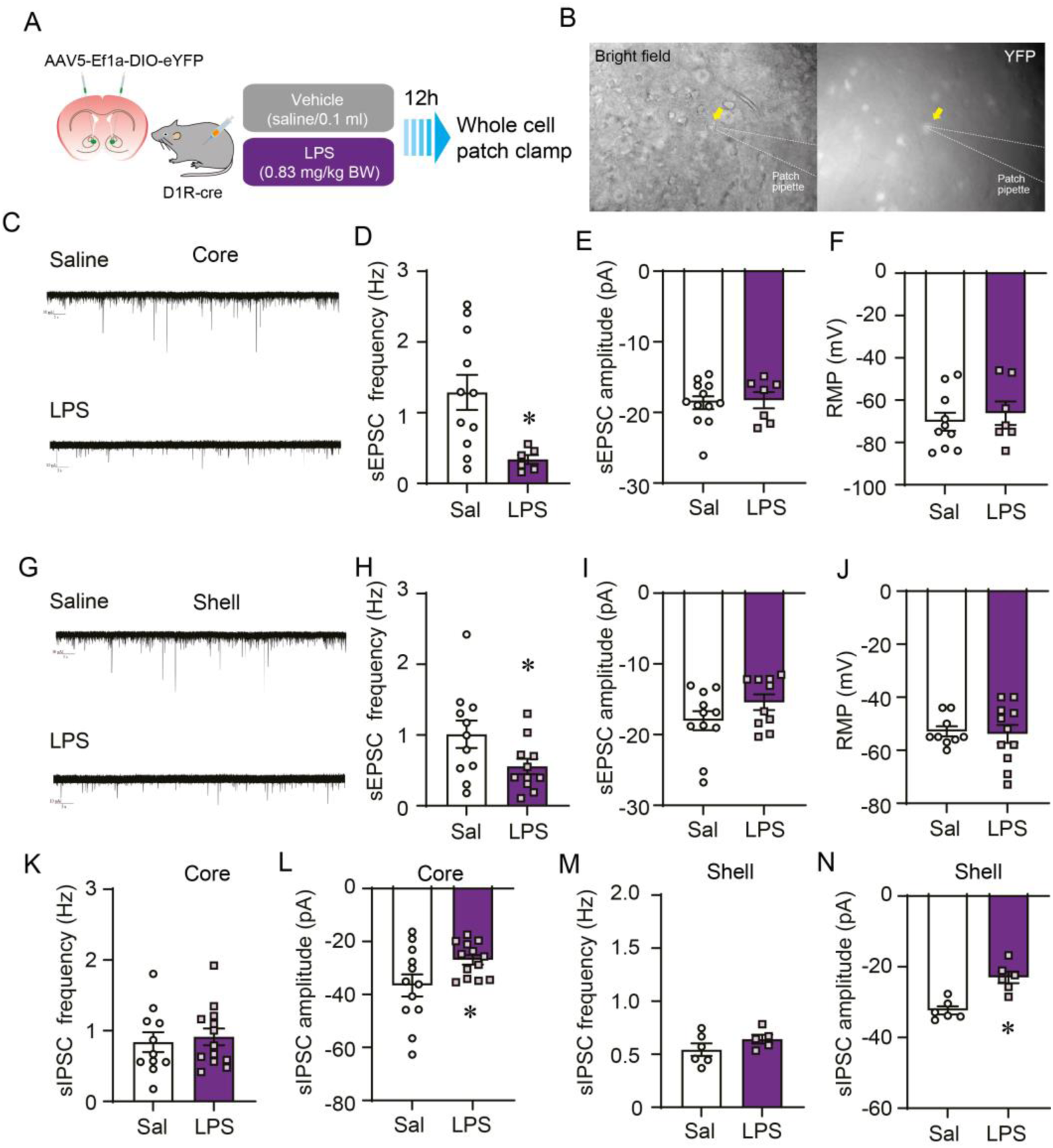
Reduced excitatory input onto D1-MSNs in the NAc in LPS-injected mice. (A) Whole-cell patch clamp electrophysiology was performed on LPS-injected (0.83 mg/kg, i.p.) D1R-cre mice with viral YFP expression (B) Representative images of patched cells in bright field and YFP. (C) Example EPSC traces in the NAc core. (D) Event frequency, (E) amplitude, and (F) resting membrane potentials of EPSCs of D1R neurons in the NAc core (n = 10-13 from mice for sal, n = 6-7 from mice for LPS). (G) Example EPSC traces in the NAc shell. (H) Event frequency, (I) peak amplitude, and (J) resting membrane potentials of D1R neurons of the NAc shell (n = 9-11 for sal from mice, n = 10-11 from mice for LPS. (K) Event frequency and (L) peak amplitude of IPSCs of D1R neurons in the NAc core. (M) Event frequency and (N) peak amplitude of IPSCs of D1R neurons in the NAc shell. (Data are expressed as mean ± SEM and analyzed using unpaired *t*-test or using non-parametric Mann Whitney test, *p<0.05.

### Primary MSN calcium responses are attenuated by microglia

To examine the contribution of reactive microglia to NAc neuronal activation via Ca^2+^ imaging, we used primary co-cultures containing primary neurons and microglia derived from the NAc (in consideration of microglial heterogeneity (34). We first studies Ca^2+^ dynamics in primary NAc neurons without microglia. NAc neurons did not respond to glutamate (10 µM) without dopamine pre-application (Figure S1A-C).

Primary NAc neurons were cultured with or without microglia (Figure 3A). LPS treatment to NAc cultured neurons without microglia increased Ca^2+^ levels after DA application (Figure 3B, 3E) whereas in the presence of microglia, NAc neurons lost the elevated Ca^2+^ in response to DA (Figure 3C, 3E) and glutamate (Figure 3F, 3H) with or without LPS treatment. The frequency of Ca^2+^ spikes was low in microglia-rich culture whereas LPS lead to a higher frequency (Figure 3D, 3G).

**Figure 3.**
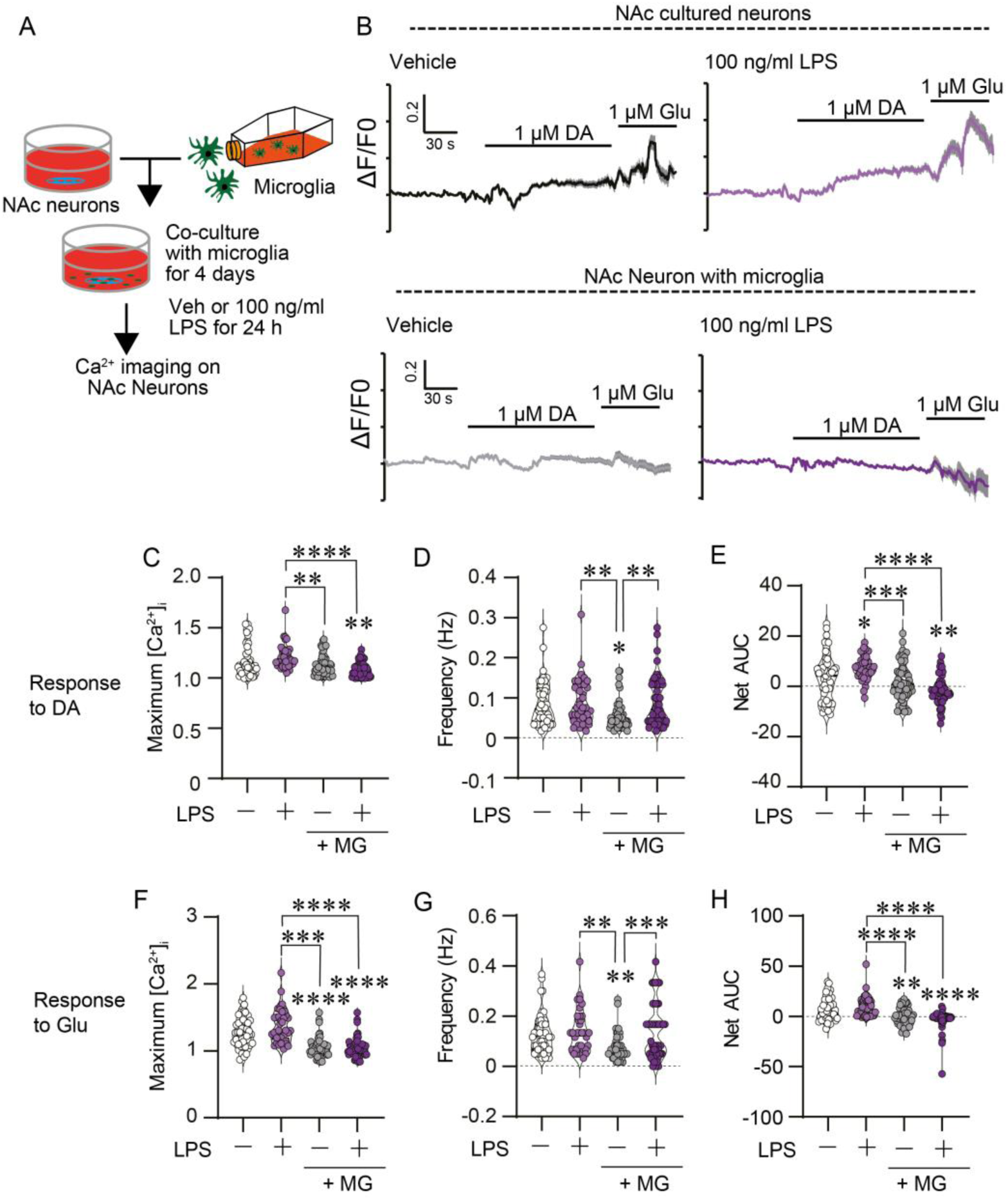
Microglial interaction abolish intracellular Ca^2+^ elevation in response to dopamine and glutamate in NAc primary cultured neurons. (A) The scheme of neuron-microglia (MG) primary co-culture for the neuronal Ca^2+^ imaging. (B) ΔF/F0 before and after 1 μM DA and 1 μM Glu with or without 100 ng/mL LPS in NAc cultured neurons and microglia rich NAc culture. (C) Maximum, (D) frequency, and (E) net AUC of [Ca^2+^] mobilization in response to DA (n = 40-54, from 3 coverslips/group. (F) Maximum, (G) frequency, and (H) net AUC of [Ca^2+^] mobilization in response to Glu (n = 35-54, from 3 coverslips/group). Data are means ± SEM. Two-way repeated measures mixed ANOVA was done followed by Bonferoni’s test. *p<0.05, **p<0.01, ***p<0.001, and ****p<0.0001.

### Increased primary microglial phagocytosis by LPS

As culture experiments demonstrated that the presence of microglial modulates NAc neuronal activity, we next explored microglial phagocytic activity in primary co-cultures of the NAc. The number (Figure 4A-B) and area (Figure 4C) of dendritic fragments (MAP2+) co-expressed in microglia (Iba1+) increased by LPS treatment.

**Figure 4.**
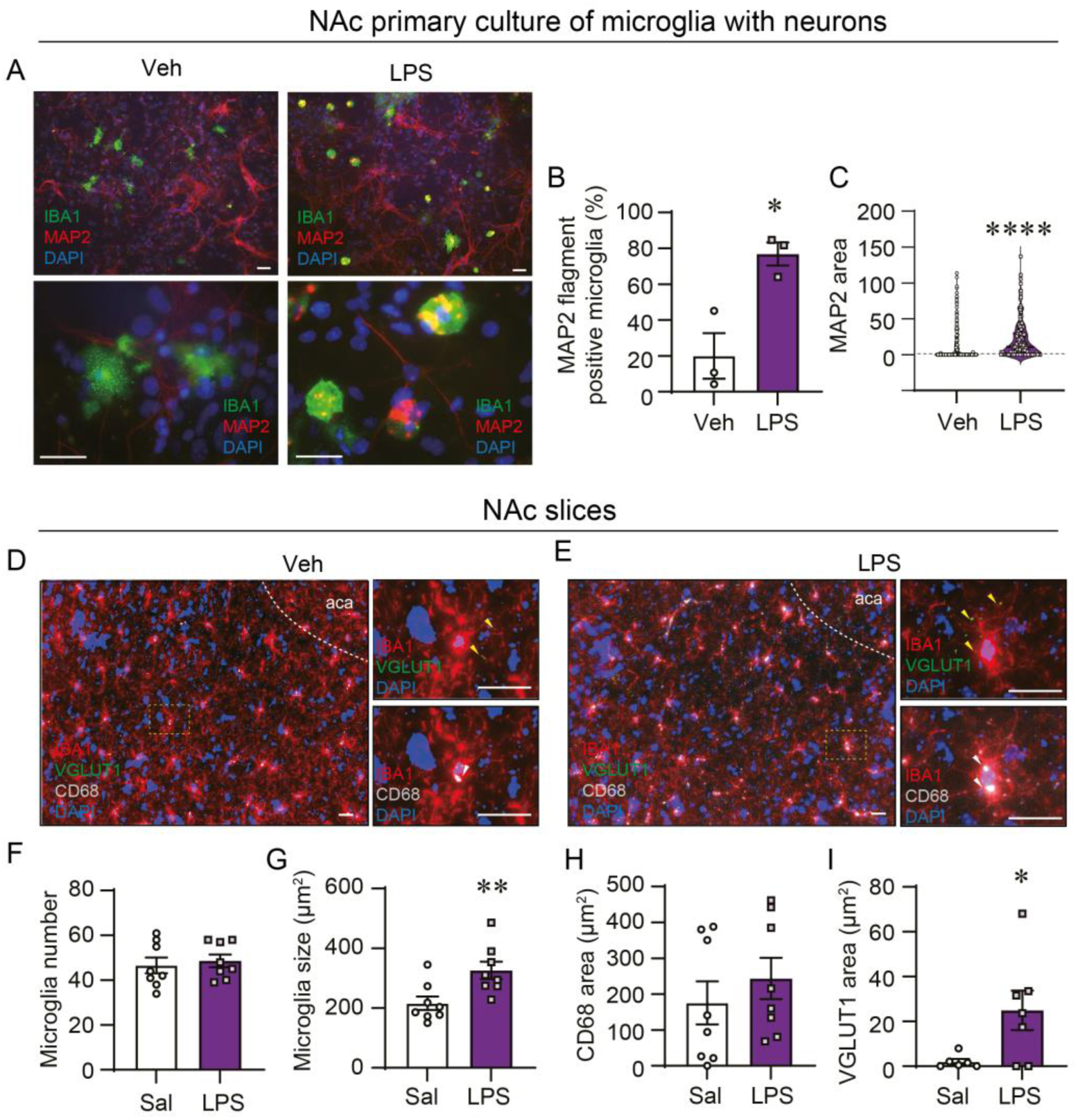
Activated microglia phagocyte neuronal process of primary cultured NAc neurons. (A) Representative images of MAP2 fragments in IBA-1 positive microglia with or without LPS treatment. Scale bar 50 µm. (B) The population of microglia containing MAP2 flagment in 3 independents primary NAc cultures. (C) Total MAP2 area in microglia (n=253 for veh, n=198 for LPS from 3 coverslips). (D) Representative images of IBA1, CD68, and VGLUT1 in the NAc from veh or LPS-treated mice. Yellow and white allow indicates VGLUT1 and CD68, respectively. Scale bar 20 µm. (F) Microglia number and (G) their size in the NAc. The area possession of (H) CD68 and (I) VGLUT1 in microglia in the NAc. Data are means ± SEM and analyzed using unpaired *t*-test. *p<0.05, **p<0.01, and ****p<0.0001.

We next sought to evaluate microglia reactivity and phagocytosis in vehicle or LPS-treated mice (Figure 4D-E). While the number of microglia in the NAc was not changed by LPS treatment (Figure 4F), NAc microglia soma size was increased by LPS treatment reflecting a more reactive (ameboid) morphology (Figure 4G). We next examined co-expression of CD68, a lysosomal glycoprotein, and Iba1 to assess microglia phagocytosis. LPS did not alter CD68 and Iba1 co-localization (Figure 4H). As CD68 upregulation in microglia is more apparent in conditions of long-term inflammation and phagocytosis of dying cells, we determined pruning of glutamatergic inputs specifically by assessing Iba1 co-localization with VGLUT1, a glutamatergic terminal marker. VGLUT1 was significantly increased in microglia in the NAc of LPS-treated mice (Figure 4I).

## DISCUSSION

The neuroinflammatory consequences of systemic immune threats can give rise to sickness-like behaviors that blunt locomotion, mood and sociability. We sought to uncover neural mechanisms contributing to such behavioral changes by inflammation by studying changes in MSN excitability, microglia activation and synaptic pruning in response to acute LPS. At a LPS dose that stimulated anxiety-like behavior and/or decreased locomotion across four behavioral tests, we found that LPS reduced the excitability of D1-MSNs in the NAc core and shell in c-Fos and patch-clamp recording experiments. That MSN activity in primary cultures was suppressed when in the presence of microglia suggests that microglia activation, phagocytosis and production of soluble mediators could be involved in reduced MSN activity. Our investigations in primary cultures, slices and *in vivo* suggest that NAc neuron-microglia interactions, notably microglia pruning of excitatory synaptic inputs, are involved in reduced D1-MSN neuronal activation and, consequently, mood and locomotor deficits.

As observed in previous studies, we found that LPS stimulated anxiety-like behavior, reduced frequency of social contacts and elicited psychomotor slowing (9, 40, 41). Behavioral indices of anxiety-like behavior were elevated in the EPM and OFT tasks 12h post LPS injection. While decreases in time spent in the light zone of the LDB test were not significant, LPS elicited reductions in the total distance traveled in the LDB. Locomotion was also blunted in the 3CT social interaction test with near-significant decreases in this parameter in the EPM and OFT to suggest that the dose we used decreases movement even following the acute sickness phase and febrile responses to LPS. LPS diminished social approach as suggested by fewer social contacts with a stranger mouse; however, LPS also produced a trend to increase contact time duration which we have observed previously with ICV cytokine administration (9). We speculate this could be due to reduced movement in general or an adaptive mechanism promoting protection during sickness.

Endotoxemia has broad effects on brain function as reflected by alterations in c-Fos immunoreactivity in multiple nuclei following LPS administration (35, 36). Glutamatergic fibers innervating the NAc originate from the ventral hippocampus, amygdala, prefrontal cortex, ventral mesencephalon and paraventricular nucleus (12, 13, 37, 38, 39). Glutamatergic transmission increases in the basolateral amygdala in LPS-treated mice, an effect blocked a serotonin reuptake inhibitor used to treat depression (40). Whereas, LPS was found to acutely elevate inhibitory tone in mPFC (41). Our results demonstrated a decrease in c-Fos positive cells in D1-MSNs in the NAc core and shell following with an increase in c-Fos in cre-negative cells that are likely largely D2-MSNs. Correspondingly, our electrophysiological findings reveal reduced excitatory input to D1-MSNs in the NAc core and shell of LPS-injected mice. We also observed a decrease in IPSC amplitude indicating a reduction in the strength of inhibitory synaptic transmission which can lead to an overall increase in neuronal excitability. The reduction in IPSC amplitude can be generated by a decrease in the number of postsynaptic GABA(A) receptors or a reduction in their clustering at synaptic sites. Whether or not such changes are caused by microglia will require further investigation. Nevertheless, reductions in c-Fos expression and reduced EPSCs suggest that acute endotoxemia dampens D1-MSNs activity in the NAc and are in agreement with a body of evidence that reduced D1-MSN activity amplifies anxiodepressive behavior (15, 42, 43). Several cell-specific approaches have uncovered D1-MSN innervation from the BLA (12, 38) and paraventricular thalamus, encoding reinforcement and aversion, respectively (13). Chemogenetic and genetic ablation of NAc D1-MSNs alters social interaction and dominance, suggesting that the balance of D1- and D2-MSNs activity contributes to sociability (42, 43). The activation of glutamatergic input into NAc D1- and D2-MSNs from BLA has been shown to reduce sociability (44). The projection of D1-MSNs onto the ventral mesencephalon and ventral pallidum are suggested to cooperatively control aversion with modulating DA balance (15). It remains unclear if and how particular D1-MSN outputs are differentially affected by systemic inflammation, however, these results support our hypothesis that inhibition of D1R neurons in the NAc core and shell are at least partially responsible for mood deficits caused by acute systemic inflammation.

Microglia are fundamental for regulating neuron health and synapse maintenance via direct and indirect cellular interactions (45). Activated microglia can secrete proinflammatory cytokines, present antigen and become phagocytic. Microglial phagocytosis is an essential process for removing cellular debris during pathological conditions (46) and restoring homeostasis in neural networks (47). Microglia have been shown to negatively modulate striatal neuronal activity in seizure models through purinergic signaling (27) and to increase NAc neuronal activity via cytokine production in conditions of cocaine behavioral sensitization (48). In our co-culture model, the presence of microglia resulted in reduced Ca^2+^ responses to DA and Glu, independent of LPS treatment. That LPS treatment was not required for reduced Ca^2+^ levels may be due to the reactive nature that non-LPS treated primary microglia largely exhibit in culture (amoeboid morphology). Nonetheless, we found that LPS treatment enhanced microglial phagocytosis of neuronal processes as shown by higher MAP2 expression in microglia in primary NAc co-cultures. CD68 is a phagocytic marker well-linked to neuroinflammatory responses in regions such as the mPFC (41), hippocampus (49, 50), and substantia nigra (51). However, CD68 levels were unchanged by LPS which may be because microglial CD68 expression is more associated with phagocytosing of degenerating or dying neurons characteristic of long-term inflammatory conditions (52). Increased VGLUT1 expression in microglia that we observed suggests that glutamatergic inputs are being engulfed by microglia, a finding that aligns with neuroinflammatory outcomes in other brain regions (28, 53, 54, 55). Together, these data suggest that pruning of excitatory synapses contribute to reduced D1-MSN activity by LPS.

In summary, systemic inflammation selectively modulates brain regions controlling sickness behavior such as resting, avoidance, anxiety, and depression (7, 8, 9, 10, 11). Microglia pruning of glutamatergic synapses that leads to reduced excitatory input onto D1-MSNs in the NAc caused by LPS-injection could be a central mechanism by which acute endotoxemia evokes sickness- and anxiety-like. Furthermore, these findings add to the growing research suggesting that the microglia play a key role in refining neuronal activity and behavior via engulfing GABAergic and/or glutamatergic synapses during inflammation (28, 53, 54, 55). As chronic inflammation is a core feature of metabolic (4), cancer (56), and psychiatric diseases, deeper investigation of microglia-neurons interactions may well have implications for understanding mood dysfunction in these conditions as well (57). Together with, the present study is a valuable insights into the neural dynamics underlying inflammation-induced anxiety and potentially inform new therapeutic approaches for mental health.

## Acknowledgements

This work was supported by research grant from Canadian Institutes of Health Research (S.F.), S.N. was supported by a fellowship from the Japan Society for the promotion of Science (JSPS) and the Uehara memorial foundation. We thank the metabolic phenotyping core of CRCHUM and Demetra Rodaros for their technical assistance.

## Author Contributions

S.N. A.K. and S.F. designed studies. S.N., A.K, S.A., S.D.N., and A.T. performed in vivo and biochemical experiments. A.K. performed electrophysiological experiments. S.N. performed primary cultured cell experiments. S.N., A.K., S.A., S.D.N., A.T., and S.F. analyzed data. A.B. and C.M.R. provided guidance in experimental design and interpretation. S.N, A.K., and S.F. wrote manuscript. All authors critically contributed to review of manuscript.

## Declaration of Interests

The authors declare no competing interests.

## Figure legends

**Supplemental Figure 1.**
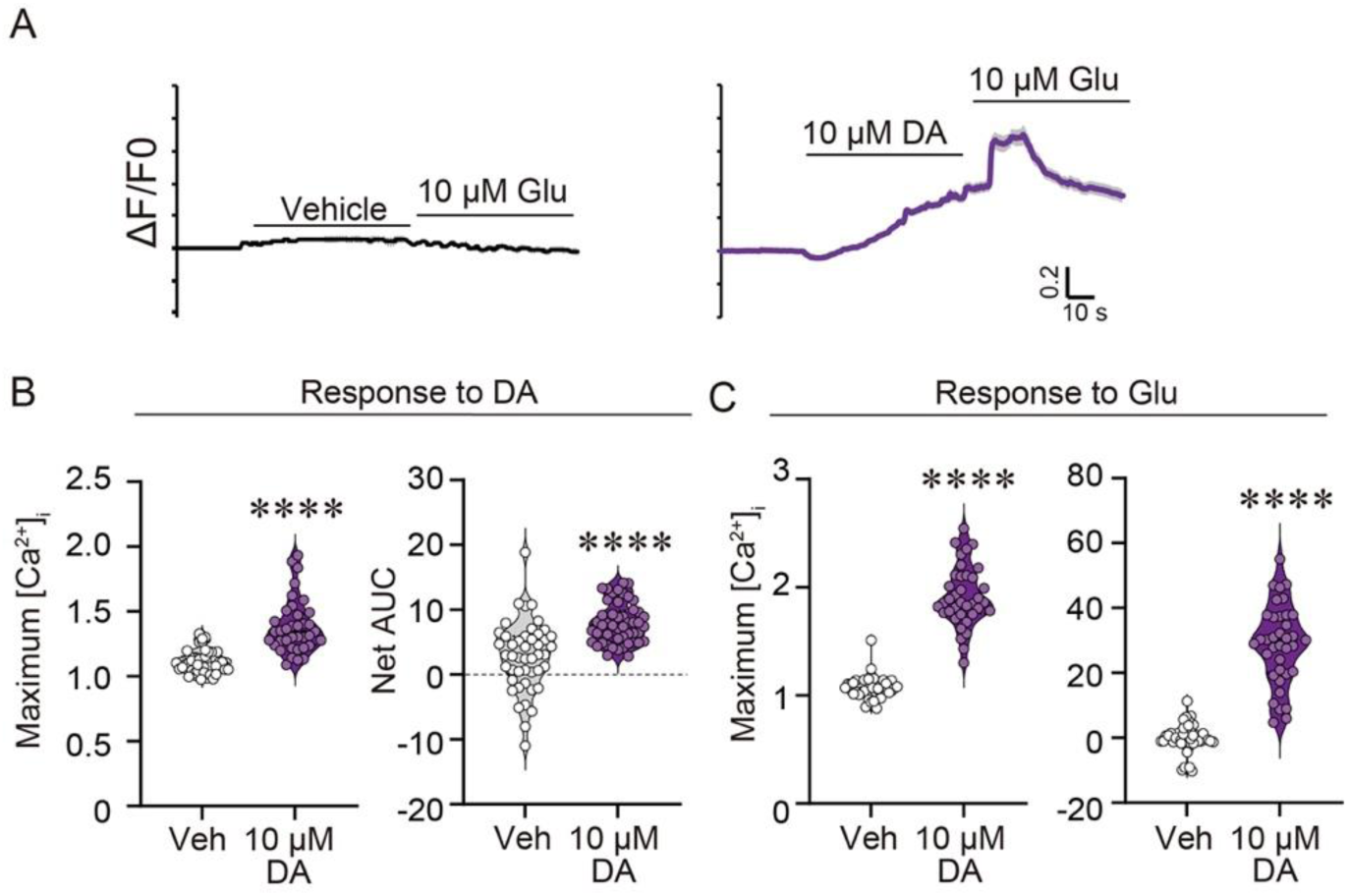
NAc neurons require dopamine stimulation on glutamate sensing. (A) The effect of dopamine on neuronal Ca^2+^ response to Glu in NAc Neurons (n = 46 for veh, n = 38 for DA from 4 coverslips) (B) Maximum and net AUC of [Ca^2+^] mobilization in response to DA. (C) Maximum and net AUC of [Ca^2+^] mobilization in response to Glu. Data are means ± SEM and analyzed using unpaired *t*-test. ****p<0.0001.

## Notes

### Competing Interest Statement

The authors have declared no competing interest.

